# Neural correlates of subsequent memory-related gaze reinstatement

**DOI:** 10.1101/2021.02.23.432536

**Authors:** Jordana S. Wynn, Zhong-Xu Liu, Jennifer D. Ryan

**Affiliations:** Department of Psychology, Harvard University, USA; Behavioral Sciences, University of Michigan-Dearborn, Michigan, USA; Rotman Research Institute at Baycrest Health Sciences, Toronto, ON, Canada; Departments of Psychology, Psychiatry, University of Toronto, Toronto, ON, Canada

## Abstract

Mounting evidence linking gaze reinstatement- the recapitulation of encoding-related gaze patterns during retrieval- to behavioral measures of memory suggests that eye movements play an important role in mnemonic processing. Yet, the nature of the gaze scanpath, including its informational content and neural correlates, has remained in question. In the present study, we examined eye movement and neural data from a recognition memory task to further elucidate the behavioral and neural bases of functional gaze reinstatement. Consistent with previous work, gaze reinstatement during retrieval of freely-viewed scene images was greater than chance and predictive of recognition memory performance. Gaze reinstatement was also associated with viewing of informationally salient image regions at encoding, suggesting that scanpaths may encode and contain high-level scene content. At the brain level, gaze reinstatement was predicted by encoding-related activity in the occipital pole and basal ganglia, neural regions associated with visual processing and oculomotor control. Finally, cross-voxel brain pattern similarity analysis revealed overlapping subsequent memory and subsequent gaze reinstatement modulation effects in the parahippocampal place area and hippocampus, in addition to the occipital pole and basal ganglia. Together, these findings suggest that encoding-related activity in brain regions associated with scene processing, oculomotor control, and memory supports the formation, and subsequent recapitulation, of functional scanpaths. More broadly, these findings lend support to Scanpath Theory’s assertion that eye movements both encode, and are themselves embedded in, mnemonic representations.

## Introduction

The human visual field is limited, requiring us to move our eyes several times a second to explore the world around us. This necessarily sequential process of selecting visual features for fixation and further processing has important implications for memory. Research using eye movement monitoring indicates that during visual exploration, fixations and saccades support the binding of salient visual features and the relations among them into coherent and lasting memory traces (e.g., Liu, Shen, Olsen, & Ryan, 2017; Liu, Rosenbaum, & Ryan, 2020; for review, see Wynn, Shen, & Ryan, 2019). Moreover, such memory traces may be stored and subsequently recapitulated as patterns of eye movements or ‘scanpaths’ at retrieval (Noton & Stark, 1971b, 1971a; for review, see Wynn et al., 2019). Specifically, when presented with a previously encoded stimulus, or a cue to retrieve a previously encoded stimulus from memory, humans (and non-human primates, see Sakon & Suzuki, 2019) spontaneously reproduce the scanpath enacted during encoding (i.e., gaze reinstatement), and this reinstatement is predictive of mnemonic performance across a variety of tasks (e.g., Damiano & Walther, 2019; Foulsham et al., 2012; Johansson & Johansson, 2013; Laeng, Bloem, D’Ascenzo, & Tommasi, 2014; Laeng & Teodorescu, 2002; Olsen, Chiew, Buchsbaum, & Ryan, 2014; Scholz, Mehlhorn, & Krems, 2016; Wynn, Olsen, Binns, Buchsbaum, & Ryan, 2018; Wynn, Ryan, & Buchsbaum, 2020; for review, see Wynn et al., 2019). While there is now considerable evidence supporting a link between gaze reinstatement (i.e., reinstatement of encoding gaze patterns during retrieval) and memory retrieval, investigations regarding the neural correlates of this effect are recent and few (see Bone et al., 2018; Ryals, Wang, Polnaszek, & Voss, 2015), and no study to date has investigated the patterns of neural activity at encoding that predict subsequent gaze reinstatement. Thus, to further elucidate the link between eye movements and memory at the neural level, the present study used concurrent eye movement monitoring and functional magnetic resonance imaging (fMRI) to investigate the neural mechanisms at encoding that predict functional gaze reinstatement (i.e., gaze reinstatement that supports mnemonic performance) at retrieval, in the vein of subsequent memory studies (e.g., Brewer, Zhao, Desmond, Glover, & Gabrieli, 1998; Wagner et al., 1998; for review, see Hannula & Duff, 2017).

Scanpaths have been proposed to at once contain, and support the retrieval of, spatiotemporal contextual information (Noton & Stark, 1971b, 1971a). According to Noton & Stark’s seminal Scanpath Theory (1971b, 1971a), on which much of the current gaze reinstatement literature is based (see Wynn et al., 2019), scanpaths consist of both image features and the fixations made to them as “an alternating sequence of sensory and motor memory traces”. Consistent with this proposal, research using eye movement monitoring and neuroimaging techniques has established an important role for eye movements in visual memory encoding (for review, see Meister & Buffalo, 2016; Ryan, Shen, & Liu, 2020). For example, at the behavioral level, recognition memory accuracy is significantly attenuated when eye movements during encoding are restricted (e.g., to a central fixation cross) as opposed to free (e.g., Damiano & Walther, 2019; Henderson, Williams, & Falk, 2005; Liu et al., 2020). At the neural level, restricting viewing to a fixed location during encoding results in attenuated activity in brain regions associated with memory and scene processing including the hippocampus (HPC) and parahippocampal place area (PPA), as well as reduced functional connectivity between these regions and other cortical regions (Liu, Rosenbaum, & Ryan, 2020). When participants are free to explore, however, the number of fixations executed is positively predictive of subsequent memory performance (e.g., Fehlmann et al., 2020; Liu et al., 2017; Loftus, 1972; Olsen et al., 2016) and of activity in the HPC (Liu et al., 2017, 2020; see also, Olsen et al., 2016) and medial temporal lobe (Fehlmann et al., 2020), suggesting that eye movements are critically involved in the accumulation and encoding of visual feature information into lasting memory traces. That the relationships between gaze fixations and recognition memory performance (e.g., Wynn, Buchsbaum, et al., 2020; see also, Chan, Chan, Lee, & Hsiao, 2018) and gaze fixations and HPC activity (Liu, Shen, Olsen, & Ryan, 2018) are reduced with age, despite an increase in the number of fixations (e.g., Firestone, Turk-Browne, & Ryan, 2007; Heisz & Ryan, 2011), further suggests that these effects extend beyond the effects of mere attention or interest.

Recent work suggests that eye movements not only play an important role in memory encoding, but also actively support memory retrieval. Consistent with Scanpath Theory, several studies have provided evidence that gaze patterns elicited during stimulus encoding are recapitulated during subsequent retrieval and are predictive of mnemonic performance (e.g., Damiano & Walther, 2019; Foulsham et al., 2012; Johansson & Johansson, 2013; Laeng et al., 2014; Laeng & Teodorescu, 2002; Olsen et al., 2014; Scholz et al., 2016; Wynn et al., 2018, 2020, for review, see 2019). In addition to advancing a functional role for eye movements in memory retrieval, this literature has raised intriguing questions regarding the nature of the scanpath and its role in memory. For example, how are scanpaths created, and what information do they contain? To answer these questions, it is necessary not only to relate eye movement and behavioral patterns, as prior research has done, but also, and perhaps more critically, to relate eye movement and neural patterns. Yet, only two studies, to our knowledge, have directly investigated the neural correlates of gaze reinstatement, with both focusing on retrieval-related activity patterns. In the first of these studies, Ryals et al. (2015) demonstrated that trial-level variability in gaze similarity (between previously viewed scenes and novel scenes with similar feature configurations), was associated with activity in the right HPC. Extending this work, Bone and colleagues (2018) observed that gaze reinstatement (i.e., similarity between participant- and image-specific gaze patterns during encoding and subsequent visualization) was positively correlated with whole-brain neural reinstatement (i.e., similarity between image-specific patterns of brain activity evoked during encoding and subsequent visualization) during a visual imagery task. Considered together, these two studies provide evidence that functional gaze reinstatement is related to neural activity patterns typically associated with memory retrieval, suggesting a common mechanism.

While there is now some evidence that mnemonic retrieval processes support gaze reinstatement at the neural level, the relationship between gaze reinstatement and encoding-related neural activity has yet to be investigated. Accordingly, the present study used the data from Liu et al. (2020) to elucidate the encoding mechanisms that support the formation and subsequent recapitulation of functional scanpaths. Participants encoded intact and scrambled scenes under free or fixed (restricted) viewing conditions (in the scanner) and subsequently completed a recognition memory task with old (i.e., encoded) and new (i.e., lure) images (outside of the scanner). Previous analysis of this data revealed that when compared to free viewing, restricting eye movements reduced activity in the hippocampus, connectivity between the hippocampus and other visual and memory regions, and ultimately reduced subsequent memory performance (Liu et al., 2020). These findings critically suggest that eye movements and memory encoding are linked at both the behavioral and neural level. Here, we extend this work further by investigating the extent to which the patterns of eye movements, or scanpaths, that are created at encoding are reinstated at retrieval to support memory performance, and by investigating the neural activity at encoding that predicts the subsequent reinstatement of scanpaths at retrieval.

To this end, we first computed the spatial similarity between encoding and retrieval scanpaths (containing information about fixation location and duration) and used this measure to predict recognition memory accuracy. Based on prior evidence of functional gaze reinstatement, we predicted that gaze reinstatement would be both greater than chance and positively correlated with recognition of old images. To further interrogate the nature of information represented in the scanpath, we additionally correlated gaze reinstatement with measures of visual (i.e., stimulus-driven; bottom-up) and informational (i.e., participant-driven; bottom-up and top-down) saliency. Given that prior work has revealed a significant role for top-down features (e.g., meaning, Henderson & Hayes, 2018; scene content, O’Connell & Walther, 2015) in guiding eye movements, above and beyond bottom-up image features (e.g., luminance, contrast, Itti & Koch, 2000), we hypothesized that gaze reinstatement would be related particularly to the viewing of informationally salient image regions. Finally, to uncover the neural correlates of functional gaze reinstatement, we analyzed neural activity patterns at encoding, both across the whole-brain and in memory-related regions of interest (i.e., HPC, PPA, see Liu et al., 2020), to identify brain regions that (1) predicted subsequent gaze reinstatement at retrieval, and (2) showed overlapping subsequent gaze reinstatement and subsequent memory effects. Given that previous work has linked gaze scanpaths, as a critical component of mnemonic representations, to successful encoding and retrieval, we hypothesized that functional gaze reinstatement would be supported by encoding-related neural activity in brain regions associated with visual processing (i.e., ventral visual stream regions) and memory (i.e., medial temporal lobe regions). By linking the neural correlates and behavioral outcomes of gaze reinstatement, the present study provides novel evidence in support of Noton & Stark’s assertion that scanpaths both serve to encode, and are themselves encoded into memory, allowing them to facilitate retrieval via recapitulation and reactivation of informationally salient image features.

## Methods

### Participants

Participants were 36 young adults (22 female) aged 18-35 (*M* = 23.58, *SD* = 4.17) with normal or corrected-to-normal vision and no history of neurological or psychiatric disorders. All participants were recruited from the University of Toronto and surrounding Toronto area community and were given monetary compensation for their participation in the study. All participants provided written informed consent in accordance with the Research Ethic Board at the Rotman Research Institute at Baycrest Health Sciences.

### Stimuli

Stimuli consisted of 864, 500 × 500-pixel, colored images, made up of 24 images of each of 36 semantic scene categories (e.g., living room, arena, warehouse, etc.), varying along the feature dimensions of size and clutter (6 levels per dimension = 36 unique feature level combinations, balanced across conditions)^1^. Within each scene category, 8 images were assigned to the free-viewing encoding condition and 8 images were assigned to the fixed-viewing encoding condition; images were randomly assigned to 8 fMRI encoding runs (36 images per run per viewing condition). The remaining 8 images in each scene category were used as novel lures at retrieval. One hundred and forty-four scene images from encoding (72 images per viewing condition from 2 randomly selected encoding runs), and 72 scene images from retrieval (2 per scene category) were scrambled using 6 levels of tile size (see Fig 1). Thus, in total, 432 intact scene images and 144 scrambled color-tile images were viewed at encoding, balanced across free- and fixed-viewing conditions, and 648 intact scene images (432 old and 216 novel lure) and 216 scrambled color-tile images (144 old and 72 novel lure) were viewed at retrieval. All images were balanced for low-level image properties (e.g., luminance, contrast)^2^ and counterbalanced across participants (for assignment to experimental/stimulus conditions).

**Fig 1.**
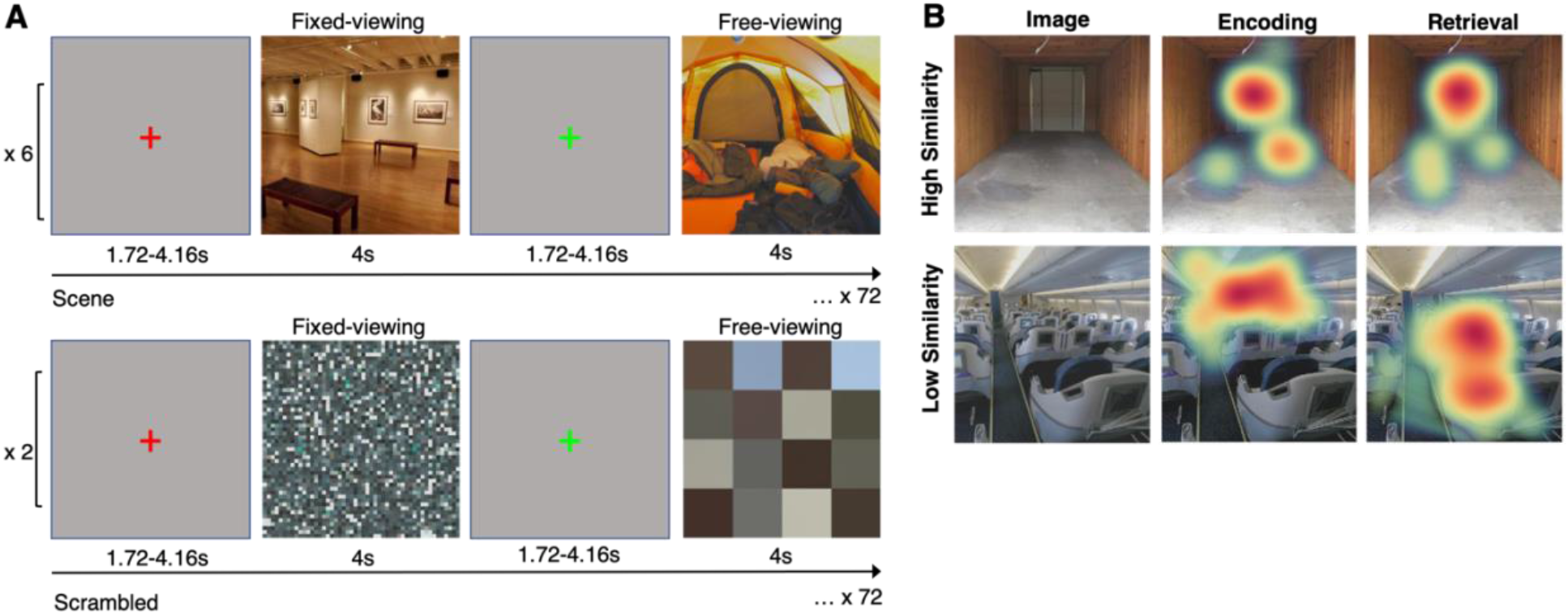
(A) Visualization of the experimental procedure for the in-scan encoding task. Prior to each trial, a green or red fixation cross was presented on the screen indicating whether participants would be required to freely view (green) or maintain fixation (red) during presentation of the upcoming image. Note that while fixations are presented centrally here, during the experiment they were presented in a random location within a 100-pixel radius around the center of the screen. Participants completed 6 runs of scenes and 2 runs of scrambled color-tile images, consisting of 72 images each. (B) Visualization of the gaze reinstatement analysis, with one example each of a high similarity and low similarity score. Reinstatement (i.e., similarity) scores reflect the spatial overlap between patterns of fixations (defined by location and duration) corresponding to the same image viewed by the same participant during encoding and retrieval, controlling for image-invariant (idiosyncratic) viewing biases (e.g., center bias).

### Procedure

#### In-scan scene encoding task

Participants completed 8 encoding runs in the scanner, 6 containing scene images and 2 containing scrambled images (run order was randomized within participant^3^), see Fig 1A. Within each run, participants viewed 72 images, half of which were studied under free-viewing instructions and half of which were studied under fixed-viewing instructions. Prior to the start of each trial, participants were presented with a fixation cross for 1.72-4.16s (exponential distribution, *M* = 2.63s) presented in a random location within a 100-pixel (1.59 degrees visual angle) radius around the center of the screen. The color of the cross indicated the viewing instructions for the following image, with free-viewing indicated by a green cross and fixed-viewing indicated by a red cross. Following presentation of the fixation cross, a scene or scrambled image appeared for 4s, during which time participants were instructed to encode as much information as possible. If the image was preceded by a red cross, participants were to maintain fixation on the location of the cross for the duration of image presentation. The length of each run was 500s, with 10s and 12.4s added to the beginning and end of the run, respectively.

### Post-scan scene recognition task

Following the encoding task, participants were given a 60-minute break before completing the retrieval task in a separate testing room. For the retrieval task, participants viewed all 576 images (432 scene images and 144 scrambled color-tile images) from the encoding task along with 288 novel lure images (216 scene images and 72 scrambled color-tile images), divided evenly into 6 blocks. Prior to the start of each trial, participants were presented with a fixation cross for 1.5s presented in a random location within a 100-pixel radius around the center of the screen (for old trials, the fixation cross was presented at the same location in which it was presented during the encoding task). Following presentation of the fixation cross, a scene or scrambled image (either old, i.e., previously viewed during encoding, or novel lure) appeared for 4s. Participants were given 3s to indicate whether the presented image was “old” or “new”, and rate their confidence in that response, via key press (*z* = high confidence “old”, *x* = low confidence “old”, *n* = high confidence “new”, *m* = low confidence “new”). To quantify recognition memory for old images, points were assigned to each response as follows: *z* = 2, *x* = 1, *m* = 0, *n* = -1.

### Eyetracking procedure

During the encoding task, monocular eye movements were recorded inside the scanner using the Eyelink 1000 MRI-compatible remote eyetracker with 1000Hz sampling rate (SR Research Ltd., Mississauga, Ontario, Canada). The eyetracker was placed inside the scanner bore (behind the participants head) and detected the pupil and corneal reflection via a mirror mounted on the head coil. During the retrieval task, monocular eye movements were recorded using the Eye-link II head mounted eyetracker with 500Hz sampling rate (SR Research Ltd., Mississauga, Ontario, Canada). To ensure successful tracking during both the encoding and retrieval tasks, nine-point calibration was performed prior to the start of the task. Online manual drift correction to the location of the upcoming fixation cross was performed between trials when necessary. As head movements were restricted in the scanner, drift correction was rarely performed. Saccades greater than 0.5° of visual angle were identified by Eyelink as eye movements having a velocity threshold of 30°/s, acceleration threshold of 8000°/s, and saccade onset threshold of 0.15°. Blinks were defined as periods in which the saccade signal was missing for 3 or more consecutive samples. All remaining samples (not identified as a saccade or blink) were classified as fixations.

### MRI protocol

As specified in Liu et al. (2020), a 3T Siemens MRI scanner with a standard 32-channel head coil was used to acquire both structural and functional images. For structural T1-weighted high-resolution MRI images, we used a standard 3D MPRAGE (magnetization-prepared rapid acquisition gradient echo) pulse sequence with170 slices and using FOV = 256 × 256 mm, 192×256 matrix, 1 mm isotropic resolution, TE/TR = 2.22/200ms, flip angle = 9 degrees, and scan time = 280 s. Functional images were obtained using T2*-weighted EPI acquisition protocol with TR = 2000ms, TE = 27ms, flip angle = 70 degrees, and FOV = 191 × 192 with 64 × 64 matrix (3 mm × 3 mm in-place resolution; slice thickness = 3.5 mm with no gap). Two hundred and fifty volumes were acquired for each run. Both structural and functional images were acquired in an oblique orientation 30° clockwise to the anterior–posterior commissure axis. Stimuli were presented with Experiment Builder (SR Research Ltd., Mississauga, Ontario, Canada) back projected to a screen (projector resolution: 1024×768) and viewed with a mirror mounted on the head coil.

### Data Analysis

#### Gaze reinstatement analysis

To quantify the spatial overlap between the gaze patterns elicited by the same participants viewing the same images during encoding and retrieval, we computed gaze reinstatement scores for each image for each participant. Specifically, we computed the Fisher z-transformed Pearson correlation between the duration-weighted fixation density (i.e., heat) map^4^ (*σ* = 80) for each image for each participant during encoding and the corresponding density map for the same image being viewed by the same participant during retrieval (‘match’ similarity) (R eyesim package: https://github.com/bbuchsbaum/eyesim), see Fig 1B. Critically, while this measure (‘match’ similarity) captures the overall similarity between encoding and retrieval gaze patterns, it is possible that such similarity reflects participant-specific (e.g., tendency to view each image from left to right), image-invariant (e.g., tendency to preferentially view the center of the screen) viewing biases. Thus, to control for idiosyncratic viewing tendencies (which were not of particular interest for the present study), we additionally computed the similarity between participant- and image-specific retrieval density maps and 50 other randomly selected encoding density maps (within participant, stimulus type, and viewing condition). The resulting 50 scores were averaged to yield a single ‘mismatch’ similarity score for each participant for each image.

Match and mismatch similarity scores were contrasted using an analysis of variance (ANOVA) with similarity value as the dependent variable and stimulus type (scene, scrambled), viewing condition (free, fixed), and similarity template (match, mismatch) as independent variables. For all subsequent analyses, gaze reinstatement was reported as the difference between match and mismatch similarity scores, thus reflecting the spatial similarity between encoding and retrieval scanpaths for the same participant viewing the same image, controlling for idiosyncratic viewing biases.

To investigate the effect of gaze reinstatement on mnemonic performance, we ran a linear mixed effects model (LMEM) on trial-level accuracy (coded for a linear effect: high confidence miss = -1, low confidence miss = 0, low confidence hit = 1, high confidence hit = 2) with fixed effects including all interactions of gaze reinstatement (match similarity – mismatch similarity; z-scored), stimulus type (scene*, scrambled), and viewing condition (free*, fixed), and random effects including random intercepts for participant and image. Backwards model comparison (*α* = .05) was used to determine the most parsimonious model (*p* values approximated with the lmerTest package, Kuznetsova, Brockhoff, & Christensen, 2017).

#### Saliency analysis

To characterize gaze patterns at encoding, and specifically, the type of information encoded into the scanpath, saliency was computed for each image using two methods. First, we used a leave one subject out cross validation procedure to generate duration-weighted informational saliency (participant data-driven) maps^5^ for each image using the aggregated fixations of all participants (excluding the participant in question) viewing that image during encoding (mean number of fixations per image = 204, aggregated from all included participants). Second, we used the Saliency Toolbox (Walther & Koch, 2006) to generate visual saliency maps by producing 204 pseudo-fixations for each image based on low-level image properties including color, intensity, and orientation. Critically, whereas the stimulus (Saliency Toolbox) -driven saliency map takes into account primarily bottom-up stimulus features (e.g., luminance, contrast), the participant data-driven saliency map takes into account any features (bottom-up or top-down) that might attract viewing for any reason (e.g., semantic meaning, memory). To quantify the extent to which individual gaze patterns during encoding were guided by salient bottom-up and top-down features, participant- and image-specific encoding gaze patterns were correlated with both the informational (participant data-driven) and visual (stimulus (Saliency Toolbox)-driven) saliency maps in the same manner as the gaze reinstatement analysis described above. This analysis yielded two scores per participant per image reflecting the extent to which fixations at encoding were guided by high-level image features (i.e., informational saliency; based on the data-driven saliency map) and low-level image features (i.e., visual saliency; based on the stimulus-driven saliency map).

To investigate the relationship between encoding gaze patterns and gaze reinstatement, we ran a LMEM on gaze reinstatement with visual and informational saliency scores (z-scored) as predictors. To compare the strength of each saliency score in predicting gaze reinstatement, saliency scores were dummy coded (visual saliency = 0, informational saliency = 1). Random intercepts for participant and image were also included in the model.

#### fMRI data preprocessing

The fMRI preprocessing procedure was previously reported in Liu et al. (2020); for completeness, it is re-presented here. MRI images were processed using SPM 12 (Statistical Parametric Mapping, Welcome Trust Center for Neuroimaging, University College London, UK; https://www.fil.ion.ucl.ac.uk/spm/software/spm12/ Version: 7487) in the Matlab environment (The MathWorks, Natick, USA). Following the standard SPM 12 preprocessing procedure, slice timing was first corrected using *sinc* interpolation with the midpoint slice as the reference slice. Then, all functional images were aligned using a six-parameter linear transformation. Next, for each participant, functional image movement parameters obtained from the alignment procedure, as well as the global signal intensity of these images, were checked manually using the freely available toolbox ART (http://www.nitrc.org/projects/artifact_detect/) to detect volumes with excessive movement and abrupt signal changes. Volumes indicated as outliers by ART default criteria were excluded later from statistical analyses. Anatomical images were co-registered to the aligned functional images and segmented into white matter (WM), gray matter (GM), cerebrospinal fluid (CSF), skull/bones, and soft tissues using SPM 12 default 6-tissue probability maps. These segmented images were then used to calculate the transformation parameters mapping from the individuals’ native space to the MNI template space. The resulting transformation parameters were used to transform all functional and structural images to the MNI template. For each participant, the quality of co-registration and normalization was checked manually and confirmed by two research assistants. The functional images were finally resampled at 2×2×2 mm resolution and smoothed using a Gaussian kernel with an FWHM of 6 mm. The first five fMRI volumes from each run were discarded to allow the magnetization to stabilize to a steady state, resulting in 245 volumes in each run.

#### fMRI analysis

##### Parametric modulation analysis

To interrogate our main research question, that is, which brain regions’ activity during encoding was associated with subsequent gaze reinstatement, we conducted a parametric modulation analysis in SPM 12. Specifically, we first added the condition mean activation regressors for the *free-* and *fixed-viewing* conditions, by convolving the onset of trials of each condition with the canonical hemodynamic function (HRF) in SPM 12. We then added the trial-wise gaze reinstatement measure as our interested linear modulator, which was also convolved with the HRF. We also added motion parameters, as detailed in Liu et al. (2020), as regressors of no interest. Default high-pass filters with a cut-off of 128s and a first-order autoregressive model AR(1) were also applied.

Using this design matrix, we first estimated the modulation effect of gaze reinstatement at the individual level. These beta estimates, averaged across all scene runs, were then carried to the group level analyses in which within-subject *t* tests were used to examine which brain regions showed stronger activity when greater gaze reinstatement was observed. For this analysis, we primarily focused on the *free-viewing* scene condition as this is the condition in which the gaze reinstatement measure is most meaningful (since participants were allowed to freely move their eyes). In this analysis, the HPC and PPA served as our *a priori* ROIs (see Supplementary figure S1 in Liu et al., 2020). As specified in Liu et al. (2020), the HPC ROI for each participant was obtained using Freesurfer *recon-all* function, version 6.0 (http://surfer.nmr.mgh.harvard.edu.myaccess.library.utoronto.ca; Fischl, 2012). The PPA ROIs were obtained using the *scene vs. scrambled color tile* picture contrast. The MNI coordinates for the peak activation of the PPA were [32, -34, -18] for the right PPA and [-24, -46, -12] for the left PPA. The left and right PPA ROI contained 293 and 454 voxels, respectively.

To explore whether other brain regions showed gaze reinstatement modulation effects, in addition to the ROI analysis, we also obtained voxel-wise whole brain results. As an exploratory analysis, we used a relatively lenient threshold of *p* = .005 with 10 voxel extension (no correction), which can also facilitate future meta-analyses (Lieberman & Cunningham, 2009).

##### Brain activation pattern similarity between parametric modulation of gaze reinstatement and subsequent memory

To understand the extent to which there was similar modulation of brain activity by gaze reinstatement and by subsequent memory, we calculated cross-voxel brain activation pattern similarity between the two parametric modulation effects. This analysis allowed us to test whether the brain activity associated with the two behavioral variables (i.e., trial-wise gaze reinstatement and subsequent memory) share a similar pattern. First, we obtained subsequent memory modulation effects as detailed in Liu et al. (2020). Specifically, in this subsequent memory effect analysis, we coded subsequent recognition memory for each encoding trial based on participants’ hit/miss response and confidence (correct recognition with high confidence = 2, correction recognition with low confidence = 1, missed recognition with low confidence = 0, missed recognition with high confidence = -1). We then used this measure as a linear parametric modulator to find brain regions that showed subsequent a memory effect, that is, stronger activation when trials were subsequently better remembered. We averaged the subsequent memory effect estimates across runs for each participant. We then extracted unthresholded voxel-by-voxel subsequent memory effects and gaze reinstatement effects (i.e., estimated betas) for the HPC and PPA, separately. These beta values were then vectorized and Pearson correlations were calculated between the two vectors of the two modulation effects for each ROI. Finally, these Pearson correlations were Fisher’s Z transformed to reflect the cross-voxel pattern similarity between the subsequent memory effect and the gaze reinstatement modulation effect.

Although we mainly focused on the brain activation pattern similarity between the two modulation effects in the scene *free-viewing* condition, we also obtained the same measure for the scene *fixed-viewing* condition to provide a control condition. If the brain activation pattern modulated by the gaze reinstatement measure is related to memory processing in the scene *free-viewing* condition, it should show larger-than-zero pattern similarity with the subsequent memory effects, which should also be greater than those in the scene *fixed-viewing* condition. Therefore, at the group level, we used one-sample *t* tests to examine whether the similarity Z scores in the scene *free-viewing* condition were larger than zero and used paired *t* test to compare the similarity scores against those in the *fixed-viewing* scene condition.

In addition to the ROI brain activation pattern similarity, we also examined brain activation similarity between subsequent memory and gaze reinstatement for the whole brain in each voxel using a search-light analysis (The Decoding Toolbox v3.997; Hebart, Görgen, Haynes, & Dubois, 2015). Specifically, for each voxel, we applied an 8-mm spheric search-light to calculate the across-voxel (voxels included in this search-light) brain activation pattern similarity between the subsequent memory effect and gaze reinstatement modulation effect, using the same procedure detailed above for the ROI analysis. We first generated the brain activation similarity Z score images for the scene *free-* and *fixed-viewing* conditions separately for each participant. At the group level, the individual participants’ brain activation similarity Z score images were tested against zero for the scene *free-viewing* condition and compared to the similarity images in the scene *fixed-viewing* condition using paired *t* tests. For this whole-brain voxel-wise analysis, we used a threshold of *p* = .005 with 10 voxel extension (uncorrected; see Lieberman & Cunningham, 2009).

## Results

### Behavioral Results

Results of the ANOVA on recognition memory performance are reported in Liu et al., 2020. In short, a significant interaction of stimulus type by viewing condition indicated that recognition memory was significantly higher in the free-viewing condition than in the fixed-viewing condition, for scene images only, and for scenes relative to scrambled images, for free-viewing only (see Fig 2E in Liu et al., 2020).

**Fig 2.**
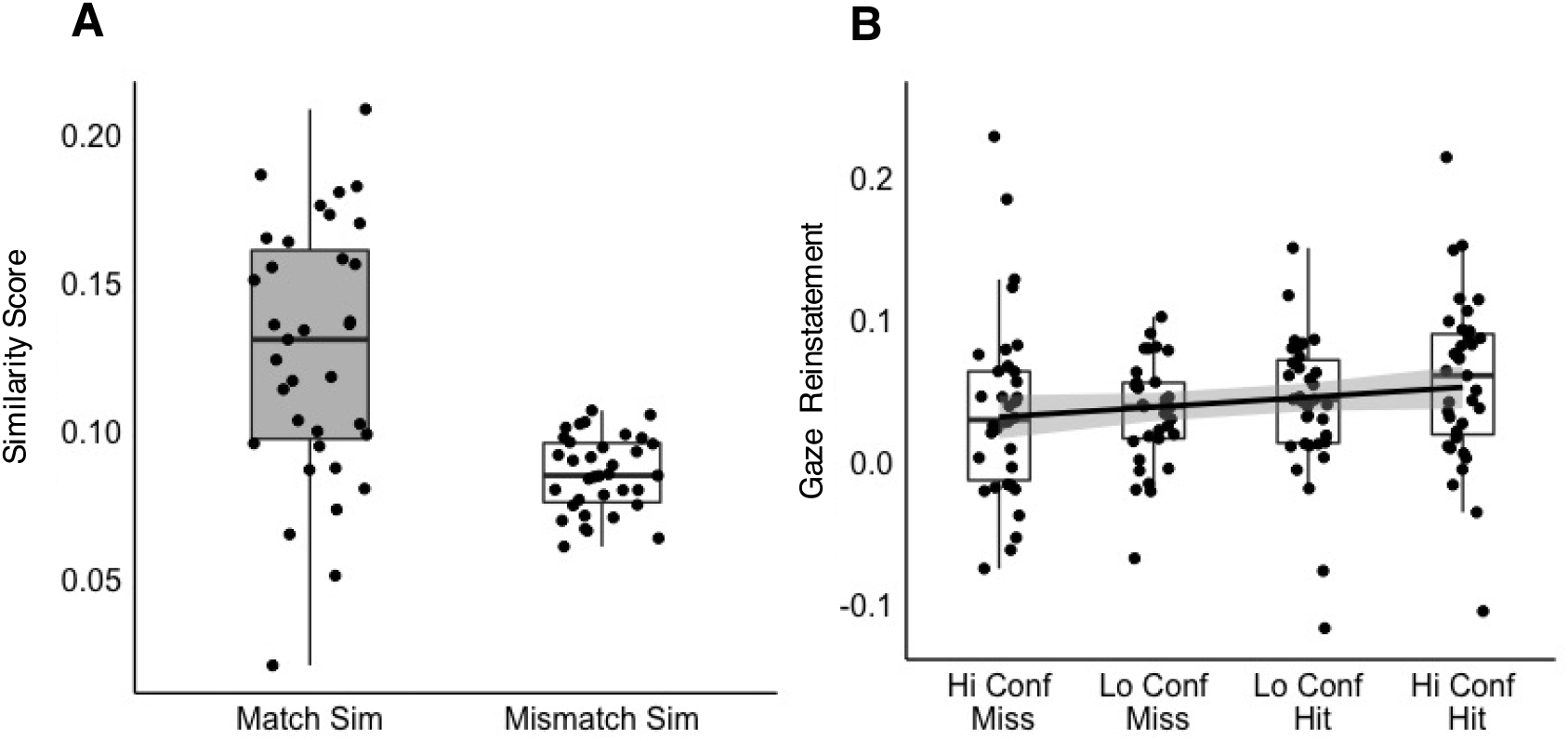
Visualization of gaze reinstatement effect for free viewing of scenes. (A) Match similarity vs. mismatch similarity scores. (B) Gaze reinstatement (match similarity-mismatch similarity) scores as a function of recognition memory accuracy.

### Eye Movement Results

To determine whether gaze reinstatement was significantly greater than chance, we ran an ANOVA with similarity value as the dependent variable and stimulus type (scene, scrambled), viewing condition (free, fixed), and similarity template (match, mismatch) as independent variables. If individual retrieval gaze patterns are indeed image specific, they should be more similar to the gaze pattern for the same image viewed at encoding (match) than for other images within the same participant, image category, and condition (mismatch). Results of the ANOVA revealed a significant 3-way interaction of similarity template, stimulus type, and viewing condition [*F* (1,34) = 7.09, *p* = .012, *η_p_* ^2^= .17]. *Post-hoc* tests of the difference in mean match and mismatch similarity scores indicated that match similarity was significantly greater than mismatch similarity in all conditions and categories [fixed scene: *t* (69.7) = 2.12, *p* = .037; fixed scrambled: *t* (69.7) = 3.60, *p* = .001; free scene: *t* (69.7) = 6.22, *p* < .001, see Fig 2A; free scrambled: *t* (69.7) = 4.583, *p* < .001].

To explore the relationship between gaze reinstatement and mnemonic performance, we ran a LMEM on trial-level accuracy with interactions of gaze reinstatement (match similarity – mismatch similarity), stimulus type (scene*, scrambled), and viewing condition (free*, fixed) as fixed effects, and participant and image as random effects. Results of the final best fit model indicated that accuracy was significantly greater for scenes relative to scrambled images (*β* = - 0.24, *SE* = 0.03, *t* = -8.19, *p* < .001, see Fig 2A), and this effect was significantly attenuated for fixed-viewing (stimulus type x viewing condition: *β* = 0.17, *SE* = 0.03, *t* = 5.10, *p* < .001). Accuracy was also significantly greater for free- relative to fixed-viewing (*β* = -0.17, *SE* = 0.16, *t* = -10.56, *p* < .001), and this effect was significantly attenuated for scrambled images (see stimulus type x viewing condition). Finally, the model revealed a significant positive effect of gaze reinstatement on accuracy (*β* = 0.06, *SE* = 0.01, *t* = 5.28, *p* < .001) for free-viewing scenes, and this effect was significantly attenuated for fixed-viewing (gaze reinstatement x viewing condition: *β* = -0.04, *SE* = 0.14, *t* = -2.82, *p* = .005) and for scrambled images (gaze reinstatement x stimulus type: *β* = -0.06, *SE* = 0.16, *t* = -3.53, *p* < .001). The addition of number of gaze fixations to the model significantly improved the model fit (*χ^2^* = 15.52, *p* < .001; see also, Liu et al., 2020), but importantly did not abolish the effect of gaze reinstatement. Furthermore, a correlation of mean gaze reinstatement scores and mean cumulative encoding gaze fixations was non-significant (*r* = .049, *p* = .79), suggesting that these effects were independent.

To determine whether gaze reinstatement (i.e., the extent to which encoding gaze patterns were recapitulated at retrieval) was related to gaze patterns (i.e., the types of information viewed) at encoding, we derived two measures to capture the extent to which individual gaze patterns at encoding reflected ‘salient’ image regions. Given that ‘saliency’ can be defined by both bottom-up (e.g., bright) and top-down (e.g., meaningful) image features, with the latter generally outperforming the former in predictive models (e.g., Henderson & Hayes, 2018; O’Connell & Walther, 2015), we computed two saliency maps for each image using the Saliency Toolbox (visual saliency map, reflecting bottom-up stimulus features) and aggregated participant data (informational saliency map, reflecting bottom-up and top-down features). Gaze patterns for each participant for each image were compared to both the visual and informational saliency maps, yielding two saliency scores. To probe the relationship between encoding gaze patterns and subsequent gaze reinstatement, we ran a LMEM on gaze reinstatement with saliency scores (visual*, informational) as fixed effects and participant and image as random effects. Results of the model revealed a significant effect of saliency on controlled gaze reinstatement (*β* = 0.10, *SE* = 0.01, *t* = 9.36, *p* < .001), indicating that similarity of individual encoding gaze patterns to the visual saliency map predicted subsequent gaze reinstatement at retrieval. Notably, the saliency effect was significantly increased when the informational saliency map was used in place of the visual saliency map (*β* = 0.06, *SE* = 0.01, *t* = 5.01, *p* < .001), further indicating that gaze reinstatement is best predicted by encoding gaze patterns that prioritize ‘salient’ image regions, being regions high in bottom-up and/or top-down informational content.

### fMRI Results

To answer our main research question regarding the neural activity patterns at encoding that predict subsequent gaze reinstatement (at retrieval), we first examined the brain regions in which activations during encoding were modulated by trial-wise subsequent gaze reinstatement scores (i.e., brain regions that showed stronger activation for trials with higher subsequent gaze reinstatement). Our ROI analyses did not yield significant effects for either the HPC or PPA (*t* = -.31 ∼ 1.13, *p* = .76 ∼ .26. Figure 3A). However, as evidenced by the whole-brain voxel-wise results (Fig 3B), the occipital poles bilaterally showed a parametric modulation by subsequent gaze reinstatement at *p* = .005, with 10 voxel extension (no correction). Two clusters in the basal ganglia also showed effects at this threshold. All regions that showed gaze reinstatement modulation effects at this threshold are presented in Table 1.

**Fig 3.**
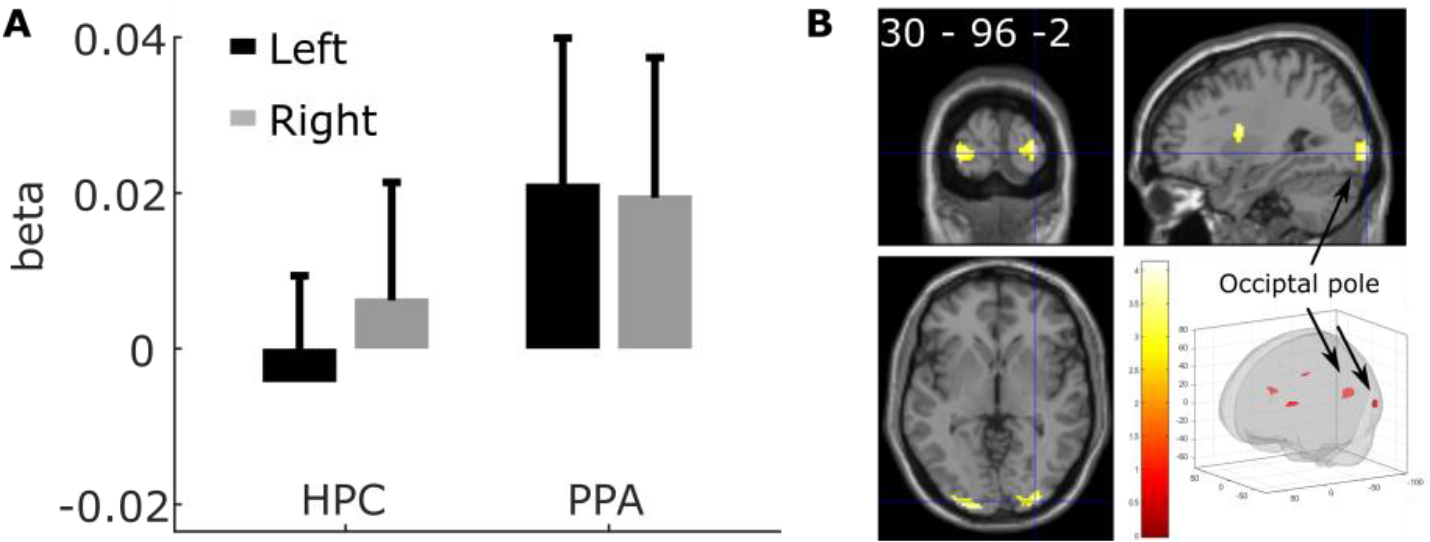
Brain activation predicted by subsequent gaze reinstatement. (A) ROI analysis revealed no significant gaze reinstatement modulation effects for HPC and PPA (all *p* > .05). (B) Voxel-wise whole brain results for gaze reinstatement modulation (thresholded at *p* = .005, 10 voxel extension, no corrections).

**Table 1.** Brain regions that positively predicted trial-wise gaze reinstatement.

| Anatomical areas | Cluster size | *t* value | *p* value | MNI coordinates |  |  |
| --- | --- | --- | --- | --- | --- | --- |
|  |  |  |  | x | y | z |
| Precentral_L | 63 | 3.983363 | 0.000164 | -30 | -4 | 42 |
| Putamen_R | 91 | 3.919645 | 0.000197 | 28 | 2 | 16 |
| Occipital_Inf_R | 180 | 3.858175 | 0.000235 | 32 | -94 | -4 |
| Occipital_Mid_L | 145 | 3.834414 | 0.000251 | -28 | -98 | 2 |
| Putamen_L | 161 | 3.815405 | 0.000265 | -26 | 12 | 12 |
| Caudate_R | 12 | 3.431526 | 0.000778 | 14 | 6 | 20 |
| Supp_Motor_Area_L | 12 | 3.112625 | 0.00184 | -10 | 16 | 50 |
| Putamen_R | 14 | 2.931331 | 0.002952 | 20 | 8 | 8 |

*Note:* All clusters survived the threshold of $p < .005$ , with 10 voxel extension, no correction. The names of the anatomical regions in the table, obtained using the AAL toolbox for SPM12, follow the automated anatomical labeling (AAL) template naming convention (Tzourio-Mazoyer et al., 2002). R/L - right/left hemisphere; Mid - middle; Inf – inferior.

As reported previously by Liu et al. (2020, see Fig 6A), both the PPA and HPC showed a parametric modulation by subsequent memory; that is, the PPA and HPC were activated more strongly for scenes that were later successfully recognized versus forgotten. Although PPA and HPC activation at the mean level were not modulated by subsequent gaze reinstatement, we investigated whether the variation across voxels within each ROI in supporting subsequent memory was similar to the variation of these voxels in supporting subsequent gaze reinstatement. Critically, this cross-voxel brain modulation pattern similarity analysis can reveal whether the pattern of activation of voxels in an ROI contains shared information, or supports the overlap, between subsequent memory and subsequent gaze reinstatement effects. Results of this analysis revealed significant pattern similarity between the two modulation effects, subsequent memory and gaze reinstatement, in both the right PPA and right HPC, *t* = 2.37 and 3.31, *p* = .024 and .002, respectively. The left HPC showed a marginally significant effect, *t* = 1.88, *p* = .069, whereas the left PPA similarity effect was not significant, *t* = 1.41, *p* = .17 (Fig 4A, B).

**Fig 4.**
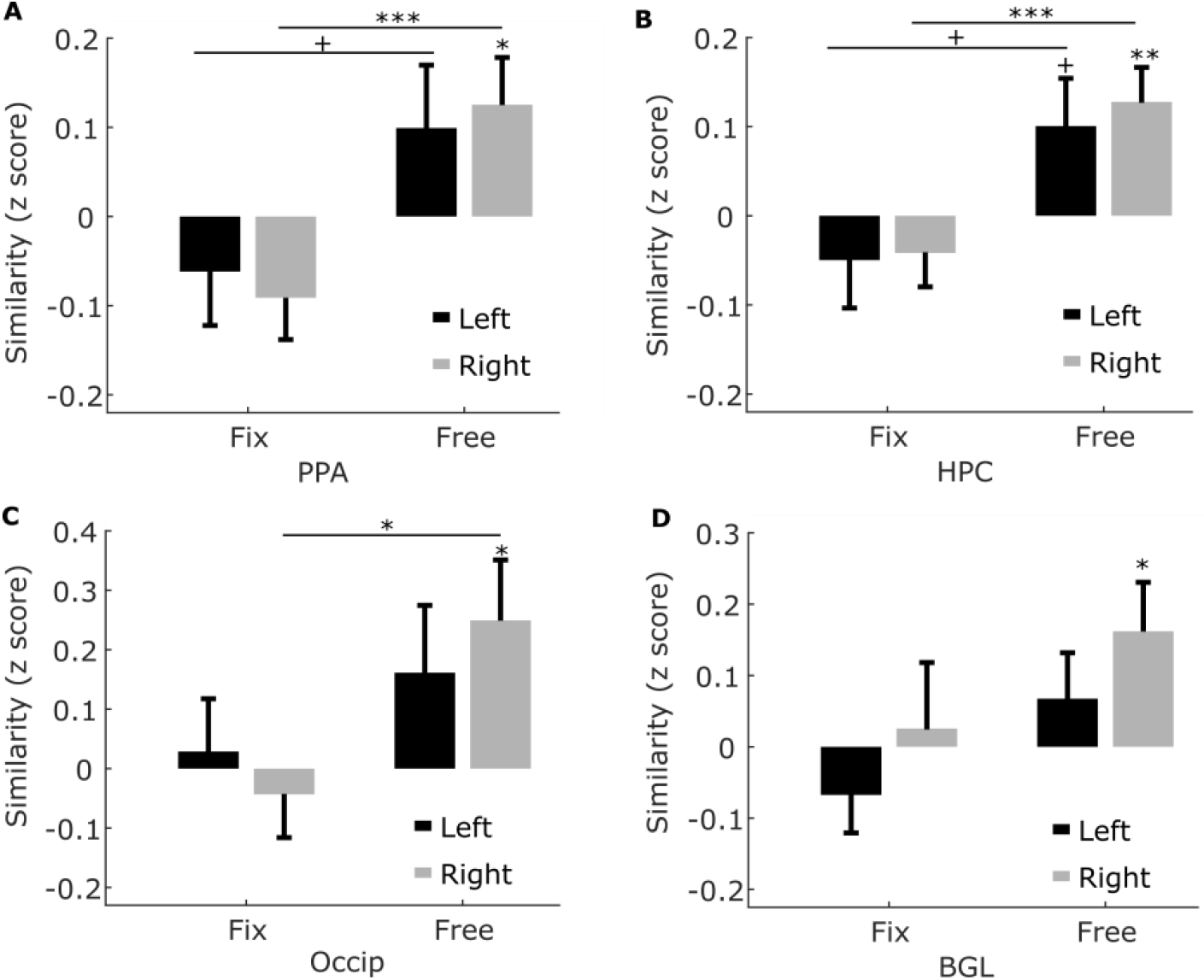
Pattern similarity between gaze reinstatement and subsequent memory modulation effects. + *p* < .09; * *p* < .05; ** *p* < .005; *** *p* < .001

**Fig 5.**
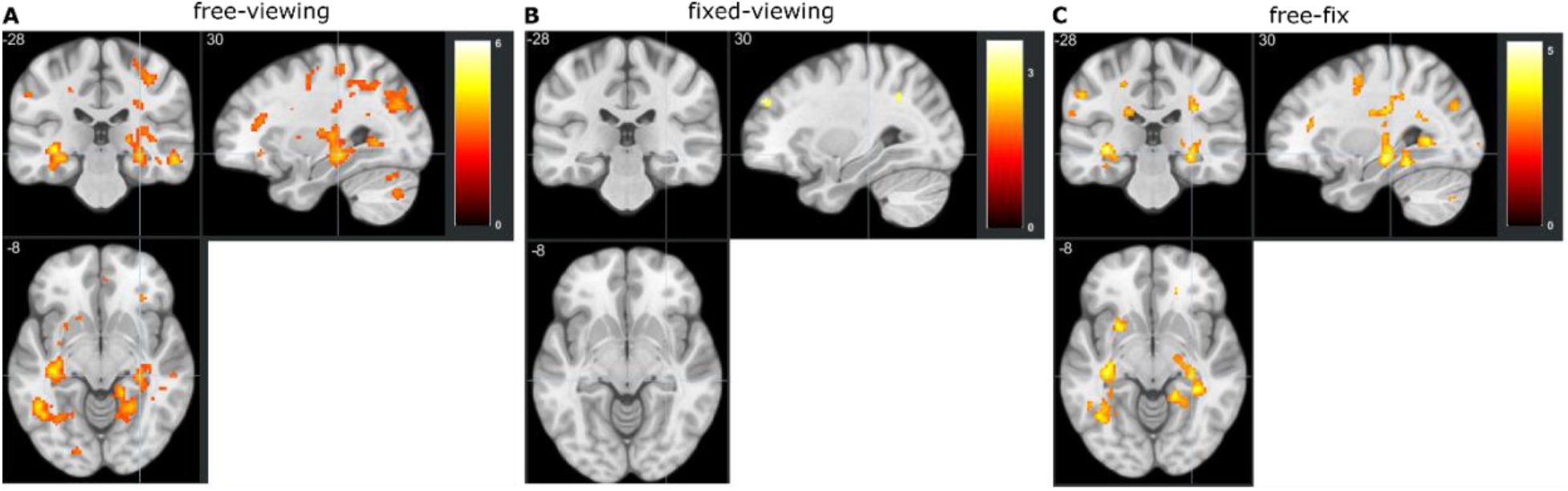
Whole-brain pattern similarity using a search light (8mm sphere) between the subsequent memory and subsequent gaze reinstatement modulation effects (threshold *p* = .005, 10 voxels extension, no corrections).

**Fig 6.**
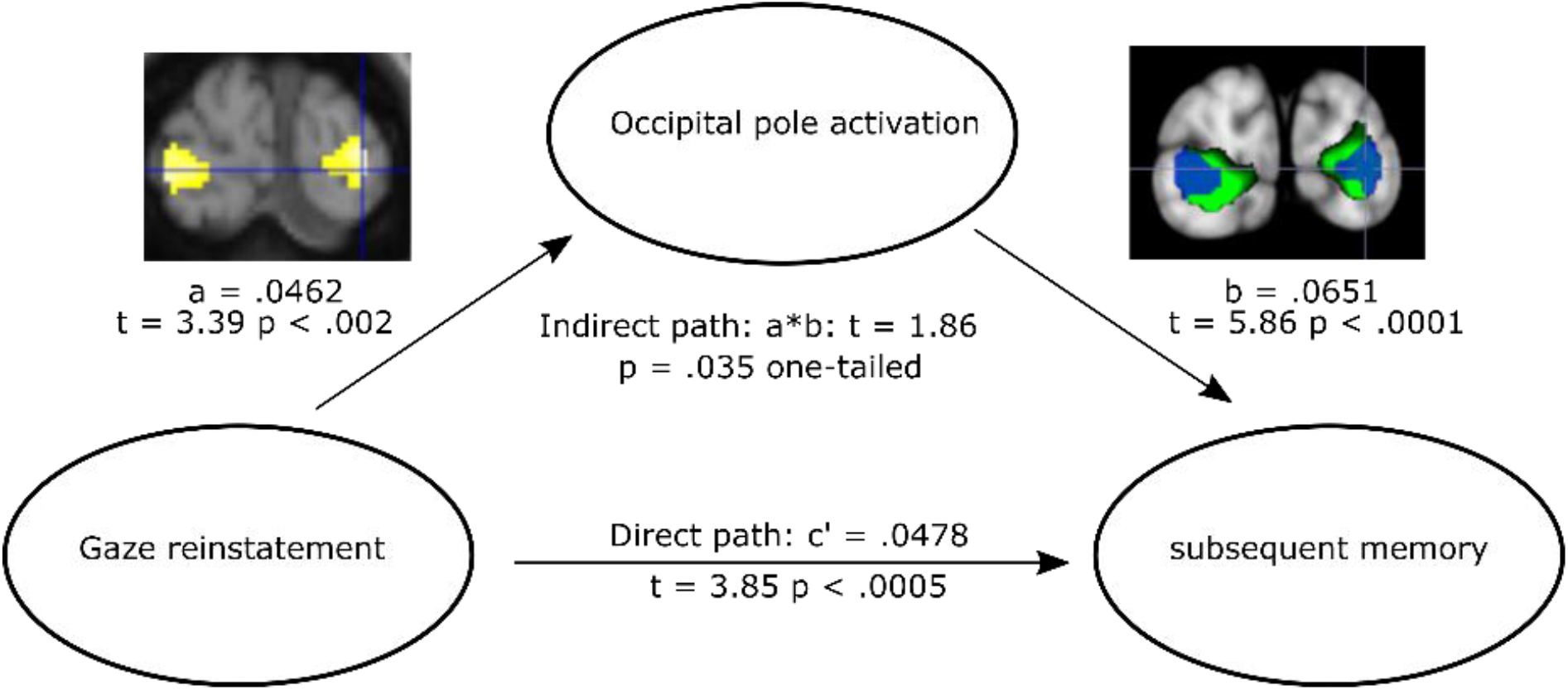
Partial mediation effect of occipital pole activation on the predictive effect of gaze reinstatement on subsequent memory. The embedded brain activation image on the left shows the occipital clusters that were modulated by subsequent gaze reinstatement. The embedded brain activation image on the right shows the overlap between the occipital clusters that were modulated by subsequent gaze reinstatement (blue) and the clusters that showed subsequent memory effects (green).

Since the occipital pole and basal ganglia regions showed stronger mean level activation for trials with greater subsequent gaze reinstatement, we also examined the pattern similarity in the voxel clusters in these two regions. Specifically, we obtained the two voxel-clusters in the basal ganglia and the occipital pole that survived the threshold of *p* = .005 (no correction) in the gaze reinstatement modulation analysis (Figure 3B) and then computed the pattern similarity scores as we did for the PPA and HPC (see above). Similar to the PPA and HPC results, the right basal ganglia and right occipital pole ROIs showed significant pattern similarity between the subsequent memory and subsequent gaze reinstatement modulation effects, *t* = 2.45 and 2.36, *p* = .02 and .024, respectively. The left ROIs did not show any significant results, *p* > .05 (Fig 4C, D).

Directly comparing the brain activation pattern similarity between the free- vs. fixed-viewing condition revealed greater brain pattern similarity in the free- vs. fixed-viewing condition for the right PPA, HPC, and occipital pole regions (*t* = 3.84, 3.55, and 2.24, *p* = .0005, .001, and .032, respectively). The left HPC and a region in the left fusiform gyrus also showed marginally significant effects (*t* = 2.02 and 1.91, *p* = .051 and .065, respectively).

As reported earlier, gaze reinstatement and memory performance were correlated at the behavioral level. Therefore, to ensure that the observed pattern similarity between the subsequent memory and gaze reinstatement modulation effects was specific to brain regions that were important for the scene encoding task, such as the ROIs tested above, and not general to all brain regions (i.e., reflecting the shared variance between the two behavioral measures at the brain level), we employed a search light method in which a sphere with radius of 8 mm was used to obtain the similarity value at each voxel of the brain. As shown in Fig 5A, not all brain regions showed the similarity effect. Instead, two large clusters in both the left and right HPC showed significant similarity between the subsequent memory and gaze reinstatement modulation effects (SPM small volume correction using HPC mask: cluster level *p*_FWE-corr_ = .004 and .012, cluster size = 248 and 172 voxels). Other regions including regions in the ventral and dorsal visual stream also showed similar patterns. These results confirm that the pattern similarity effect (i.e., the brain manifestation of the shared variance between gaze reinstatement and memory performance) occurred specifically in brain regions that are known to play key roles in visual memory encoding.

To further confirm the specificity of the pattern similarity effect, we conducted the same analysis for the fixed-viewing scene condition, which, consistent with our hypothesis, yielded no significant results in the HPC or in other ventral visual stream regions (Fig 5B). Directly contrasting the pattern similarity between the free- vs. fixed-viewing conditions confirmed that the similarity between the subsequent memory and subsequent gaze reinstatement modulation effects was specific to brain regions typically implicated in scene encoding, such as the left and right HPC (SPM small volume correction using HPC mask: cluster level *p*_FWE-corr_ = .023 and .022, cluster size = 126 and 130 voxels. Fig 5C), and specific to the free-viewing condition (Fig 5A & 5B).

Notably, the occipital poles showed stronger activation bilaterally for subsequently remembered vs. subsequently forgotten trials (embedded brain image (right) in Fig 6), and for trials with stronger subsequent gaze reinstatement (embedded brain image (left) in Fig 6). This region also showed similar cross-voxel modulation patterns for the subsequent memory and gaze reinstatement effects. We thus hypothesized that the activation of this region may mediate the relationship between gaze reinstatement and subsequent memory. To test this prediction, we conducted a mediation analysis in which we examined whether the effect of gaze reinstatement on subsequent memory could be significantly reduced when brain activity in the occipital pole, aggregated across the left and right, was entered as a mediator in the regression analysis. Specifically, for each participant, we first estimated brain activity for each scene image in each condition using the beta-series method (Rissman, et al., 2014). We then extracted the occipital pole activation corresponding to the left and right occipital pole ROIs. Next, at the individual level, we conducted a mediation analysis with the trial-wise gaze reinstatement measure as the predictor (x), occipital pole ROI activation as the mediator (m), and the subsequent memory measure as the outcome variable (y). The regression coefficient *a* (see Fig 6) was obtained when *x* was used to predict *m, b* was obtained when *m* was used to predict *y* (while controlling for *x*), and *c’* was obtained when *x* was used to predict *y* (while controlling for *m*). Finally, the coefficients *a*, *b*, and *c’* were averaged across runs for each participant and then tested at the group level using *t* tests. In line with our prediction, occipital pole activation partially mediated the prediction of gaze reinstatement on subsequent memory (indirect path: *t* = 1.87, *p* = .035, one-tailed. Fig 6.

## Discussion

The present study explored the neural correlates of functional gaze reinstatement- the recapitulation of encoding-related gaze patterns during retrieval that is significantly predictive of mnemonic performance. Consistent with Scanpath Theory (Noton & Stark, 1971b, 1971a), research using eye movement monitoring has demonstrated that the spatial overlap between encoding and retrieval gaze patterns is correlated with behavioral performance across a number of memory tasks (e.g., Damiano & Walther, 2019; Foulsham et al., 2012; Johansson & Johansson, 2013; Laeng et al., 2014; Laeng & Teodorescu, 2002; Olsen et al., 2014; Scholz et al., 2016; Wynn et al., 2018, 2020, for review, see 2019). Indeed, guided or spontaneous gaze shifts to regions viewed during encoding (i.e., gaze reinstatement) have been proposed to support memory retrieval by reactivating the spatiotemporal encoding context (Wynn et al., 2019). In line with this proposal, recent work using concurrent eyetracking and fMRI has indicated that gaze reinstatement elicits patterns of neural activity typically associated with successful memory retrieval, including HPC activity (Ryals et al., 2015) and whole-brain neural reactivation (Bone et al., 2018). Critically however, these findings do not speak to the cognitive and neural processes at encoding that support the creation of functional scanpaths. This question is directly relevant to Scanpath Theory, which contends that eye movements not only facilitate memory retrieval, but are themselves embedded in the memory trace (Noton & Stark, 1971b, 1971a). Accordingly, the present study investigated the neural regions that support the formation and subsequent recapitulation of functional scanpaths. Extending earlier findings, and lending support to Scanpath Theory, here we show for the first time that functional gaze reinstatement is correlated with encoding-related neural activity patterns in brain regions associated with sensory (visual) processing, motor (gaze) control, and memory. Importantly, these findings suggest that, like objects and the relations among them, scanpaths may be bound into memory representations, such that their recapitulation may cue, and facilitate the retrieval of, additional event elements (see Wynn et al., 2019).

Consistent with previous work, the present study found evidence of gaze reinstatement that was significantly greater than chance and significantly predictive of recognition memory accuracy when participants freely viewed repeated scenes. In addition, gaze reinstatement, (measured during free viewing of scenes at retrieval) was positively associated with encoding-related neural activity in the basal ganglia and in the occipital pole. Previous work has linked the basal ganglia to voluntary saccade control, particularly when saccades are directed towards salient or rewarding stimuli (for review, see Gottlieb, Hayhoe, Hikosaka, & Rangel, 2014), and to memory-guided attentional orienting (Goldfarb, Chun, & Phelps, 2016). Dense connections with brain regions involved in memory and oculomotor control including the HPC and frontal eye fields (FEF) (Shen, Bezgin, Selvam, McIntosh, & Ryan, 2016) make the basal ganglia ideally positioned to guide visual attention to informative (i.e., high reward probability) image regions. The occipital pole has been similarly implicated in exogenous orienting (Fernández & Carrasco, 2020) and visual processing, including visual imagery, (St-Laurent, Abdi, & Buchsbaum, 2015), partly due to the relationship between neural activity in the occipital pole and gaze measures including fixation duration (Choi & Henderson, 2015) and saccade length (Frey, Nau, & Doeller, 2020).

Notably, the occipital pole region identified in the current study was spatially distinct from the occipital place area seen in other studies (e.g., Bonner & Epstein, 2017; Dilks, Julian, Paunov, & Kanwisher, 2013; Patai & Spiers, 2017), suggesting that it may differentially contribute to scene processing, possibly by guiding visual exploration. Moreover, the identified occipital pole region did not include area V1, suggesting that unlike (the number of) gaze fixations, which modulates activity in early visual regions (Liu et al., 2017), gaze reinstatement does not directly reflect the *amount* of bottom-up visual input present at encoding. Rather, gaze reinstatement may be related more specifically to the *selective sampling* and *processing* of informative regions at encoding (see also, Fehnmann et al., 2020). Indeed, during encoding, viewing of informationally salient regions, as defined by participant data-driven saliency maps capturing both low-level and high-level image features, was significantly more predictive of subsequent gaze reinstatement than viewing of visually salient regions, as defined by stimulus (Saliency Toolbox)-driven saliency maps capturing low-level image features. The occipital pole additionally partially mediated the effect of gaze reinstatement on subsequent memory, further suggesting that this region may contribute to mnemonic processes via the formation of gaze scanpaths reflecting informationally salient image regions. Taken together with the neuroimaging results, these findings suggest that viewing, and consequentially, encoding, regions high in informational and/or rewarding content may facilitate the laying down of a scanpath that, when recapitulated, facilitates recognition via comparison of presented visual input with stored features (see Wynn et al., 2019).

To further interrogate the relationship between gaze reinstatement and memory at the neural level, we conducted a pattern similarity analysis to identify brain regions in which neural activity patterns corresponding to gaze reinstatement and those corresponding to subsequent memory covaried. Results of this analysis revealed significant overlap between the subsequent memory and subsequent gaze reinstatement effects in the occipital pole and basal ganglia (regions which showed a parametric modulation by subsequent gaze reinstatement), and in the PPA and HPC (regions which showed a parametric modulation by subsequent memory, see Liu et al., 2020). These regions may therefore be important for scene encoding (see Liu et al., 2020), in part through their role in linking the scanpath to the resulting memory representation. Specifically, parametric modulation and pattern similarity effects in the occipital pole and basal ganglia suggest that when informationally salient image features are selected for overt visual attention, those features are encoded into memory along with the fixations made to them, which are subsequently recapitulated during retrieval. Consistent with Scanpath Theory’s notion of the scanpath as a sensory-motor memory trace (Noton & Stark, 1971b, 1971a), these findings suggest that eye movements themselves may be part and parcel of the memory representation. The absence of gaze reinstatement-related activity in object- or location-specific processing regions (e.g., PPA, LOC) or low-level visual regions (e.g., V1) further suggests that reinstated scanpaths (at least in the present task) cannot be solely attributed to overlap in bottom-up visual saliency or memory for particularly salient image features. Indeed, recent work from Wang et al. (2019) indicates that simply following a face- or house-related gaze pattern (without seeing a face or house) is sufficient to elicit activity in the FFA or PPA, respectively, suggesting that visual identification is not based solely on visual features, but rather can also be supported by efferent oculomotor signals. The present findings further suggest that such signals, serving as a part of the memory representation, may be referenced and used by the HPC, similar to other mnemonic features (e.g., spatial locations, temporal order, Davachi, 2006; Yonelinas, 2013), to cue retrieval of associated elements within memory. That is, although the HPC may not be directly involved in generating or storing the scanpath (which may instead rely on visual and oculomotor regions), similar patterns of HPC activity that predict subsequent gaze reinstatement and subsequent memory suggest that the HPC may index these oculomotor programs, along with other signals, in the service of mnemonic binding and retrieval functions (e.g., relative spatial postion coding, see Connor & Knierim, 2017).

Importantly, the finding that the HPC, in particular, similarly codes for subsequent memory and subsequent gaze reinstatement is consistent with its purported role in coordinating sensory and mnemonic representations (see Knapen, 2020). Indeed, early accounts positioned the HPC as the site at which already-parsed information from cortical processors are bound into lasting memory representations (Cohen & Eichenbaum, 1993). The notion that the oculomotor effector trace is included within the HPC representation is aligned with more recent work showcasing the inherent, and reciprocal, connections between the HPC and oculomotor systems. Research using computational modeling and network analyses, for example, indicates that the HPC and FEF are both anatomically and functionally connected (Ryan, Shen, Kacollja, et al., 2019; Shen et al., 2016; for review, see Ryan, Shen, & Liu, 2019). Indeed, whereas damage to the HPC leads to impairments on several eye-movement-based measures (e.g., Hannula, Ryan, Tranel, & Cohen, 2007; Olsen et al., 2015, 2016; Ryan, Althoff, Whitlow, & Cohen, 2000), disruption of the FEF (via TMS) leads to impairments in memory recall (Wantz et al., 2016). Other work further suggests that visual and mnemonic processes share a similar reference frame, with connectivity between the HPC and V1 showing evidence of retinotopic orientation during both visual stimulation and visual imagery (Knapen, 2020; see also, Silson, Zeidman, Knapen, & Baker, 2020). That the HPC may serve as a potential ‘convergence zone’ for binding disparate event elements, including eye movements, is further supported by evidence from intracranial recordings in humans and animals suggesting that the co-ordination of eye movements with HPC theta rhythms supports memory encoding (Hoffman et al., 2013; Jutras, Fries, & Buffalo, 2013) and retrieval (Kragel et al., 2019), and by evidence of gaze-centric cells in the HPC (and entorhinal cortex, Killian, Jutras, & Buffalo, 2012; Meister & Buffalo, 2018) that respond to a particular gaze location (e.g., Chen & Naya, 2020; Rolls, Robertson, & Georges-François, 1997; for review, see Nau, Julian, & Doeller, 2018). Extending this work, the present findings suggest that gaze reinstatement and subsequent memory share similar variance in the brain and may be supported by similar HPC mechanisms. Furthermore, these findings critically suggest that reinstated gaze patterns may be recruited and used by the HPC in the service of memory retrieval.

With the present study, we provide novel evidence that encoding-related activity in the occipital pole and basal ganglia during free viewing of scenes is significantly predictive of subsequent gaze reinstatement, suggesting that scanpaths that are later recapitulated may contain important visuo-sensory and oculomotor information. Indeed, gaze reinstatement was correlated more strongly with encoding of informationally salient regions than visually salient regions, suggesting that the scanpath carries information related to high-level image content. Critically, visual, oculomotor, and mnemonic regions of interest (i.e., occipital pole, basal ganglia, PPA, HPC) showed similar patterns of activity corresponding to subsequent memory (see Liu et al., 2020) and subsequent gaze reinstatement, further supporting a common underlying neural mechanism. Lending support to Scanpath Theory, the present results suggest that gaze scanpaths, beyond scaffolding memory retrieval, are themselves embedded in the memory representation (see Cohen & Eichenbaum, 1993), similar to other elements, including spatial and temporal relations (see Davachi, 2006; Yonelinas, 2013), and may be utilized by the HPC to support memory retrieval. Given the nature of the present task, we focused here on the spatial overlap (including fixation location and duration information) between gaze patterns during encoding and retrieval, but future work could also explore how temporal order information embedded in the scanpath may similarly or differentially contribute to memory retrieval. Thus, while further research will be needed to fully elucidate the neural mechanisms supporting functional gaze reinstatement, particularly across different tasks and populations, the current findings spotlight the unique interactions between overt visual attention and memory that extend beyond behaviour to the level of the brain. Moreover, these findings speak to the importance of considering, and accounting for, effector systems, including the oculomotor system, in models of memory and cognition more broadly.

## Acknowledgments

This work was supported by a Vision: Science to Applications (VISTA) postdoctoral fellowship awarded to ZXL, funding from the Natural Sciences and Engineering Research Council of Canada awarded to JDR (RGPIN-2018-06399), and the Canadian Institutes of Health Research awarded to JDR (MOP126003).

## Footnotes

1 For further details regarding stimulus selection and feature equivalence see Liu et al. (2020).

2 To achieve luminance and contrast balance, all color RGB images were transferred to NTSC space using the built in MATLAB function rgb2ntsc.m. Then, the luminance (i.e., the NTSC Y component) and contrast (i.e., the standard deviation of luminance) were obtained for each image and the mean values were used to balance (i.e., equalize) the luminance and contrast for all images using SHINE toolbox (Willenbockek, et al., 2020). Finally, the images were transferred back to their original RGB space using the MATLAB function ntsc2rgb.m.

3 For further details regarding the randomization procedure see Liu et al. (2020).

4 For further details regarding the density map computation see Wynn, Ryan, & Buchsbaum (2020).

5 Using the same density map computation as the gaze reinstatement analysis, see Wynn, Ryan, & Buchsbaum (2020).

* Reference variable

